# Quantifying cardiac, respiratory, and slow vasomotion components of CSF motion from fMRI inflow effects

**DOI:** 10.1101/2025.06.10.658005

**Authors:** Pontus Söderström, Cecilia Björnfot, Britt M. Andersson, Jan Malm, Anders Eklund, Anders Wåhlin

## Abstract

**Purpose:** Cerebrospinal fluid (CSF) flow oscillations have emerged as a potentially important marker related to brain clearance, but their acquisition often relies on specialized imaging MRI sequences. The purpose of this work was to enable quantitative assessment of CSF flow associated with cardiac, respiratory, and slow-vasomotion cycles using widely available functional magnetic resonance imaging (fMRI) acquisitions.

**Methods:** A method was developed to translate fMRI-derived CSF inflow signals into quantitative flow rates. This approach modelled the spin-history of an oscillating ensemble of molecules. Validation was performed using phantom experiments with oscillatory flow at cardiac-, respiratory-, and slow-vasomotion-like frequencies. The method was further applied to resting-state data from 48 older adults (68–82 years; 19 women) to characterize CSF flow at the foramen magnum.

**Results:** Phantom experiments demonstrated excellent correlations between estimated and true velocities for cardiac- and respiratory-like frequencies (r = 0.94 and 0.97, respectively) and moderate correlation for a slow-vasomotion-like frequency (r = 0.58). In the population cohort, median CSF stroke volumes were 0.86 [0.61, 1.17] mL for the cardiac cycle, 0.44 [0.25, 0.94] mL for the respiratory cycle, and 0.28 [0.14, 0.45] mL for the slow-vasomotion cycle.

**Conclusion:** The proposed spin-history modeling method enabled quantitative estimation of CSF flow components using a conventional fMRI dataset and showed that the cardiac cycle dominates CSF motion at the foramen magnum.

## Introduction

Cerebrospinal fluid (CSF) flow oscillations occur in response to cardiac, respiration and low frequency vascular oscillations (1,2). Measuring this compensatory response can provide important information regarding intracranial compliance (3,4). Furthermore, CSF pulsation measurements may be informative of the physiological driving forces that can support brain clearance (5). Brain clearance, as currently understood, is performed by the glymphatic system (6,7) where CSF is suggested to follow perivascular spaces, driven by physiological mechanisms such as cardiac pulsations (8,9), respiratory-dependent pressure changes (10,11), as well as slower vascular modulations of neural origin (12,13). These physiological drivers can be studied in macroscopic CSF regions such as the 4th ventricle and the foramen magnum (14,15), and while CSF flow in the subarachnoid space is not the same as glymphatic flow, it is still believed to be related (16,17). In this context, CSF flow at the foramen magnum may be especially informative since it to a greater extent reflects the total volume of CSF displacement that happens in response to cerebrovascular volume changes, as suggested by the Monro-Kellie doctrine (18), compared to the 4th ventricle.

Functional magnetic resonance imaging (fMRI) is a widely used technique to measure neural activity non-invasively, primarily through the detection of blood oxygen level-dependent (BOLD) signals, and large databases from studies focusing on cognition in aging and dementia are already established (19–21). While the main purpose of fMRI is to detect brain activations through local changes in blood oxygenation, CSF inflow at edge slices is detectable throughout signal fluctuations. Importantly, in whole brain fMRI, the lower edge-slice typically contains CSF surrounding the spinal cord. The fMRI signal increases when unsaturated CSF enters the imaging volume, a mechanism called the inflow effect (22). By exploiting the inflow effect, researchers have indirectly measured CSF movement through signal intensity fluctuations (14). These observations have revealed that the change in global BOLD signal is coupled with the CSF flow during sleep, suggesting that CSF dynamics is linked to neural activity while sleeping. Furthermore, a significant coupling between global BOLD and CSF signal changes have also been seen during resting-state fMRI (as typically acquired in the previously mentioned large datasets), where a reduced correlation was associated with neurological disorders such as Alzheimer’s (15) and Parkinson’s disease (16). These examples, however, have isolated slower frequencies in the CSF flow spectrum, and a quantitative description regarding the importance of slow-waves, respiration and the cardiac-cycle to CSF motion is lacking.

Importantly, because of the oscillating nature of CSF flow, the inflow signal is not a direct reflection of instantaneous velocity, but rather the proportion of the fluid that has been replaced since last excitation and the spin-history. The complex dependency on spin-history adds to the challenge of translating the inflow effect to velocity, potentially obscuring true relationships between the physiological cycles and CSF oscillations. This issue becomes of particular significance when the sampling frequency is comparable to, or lower than twice the heart rate. Under such conditions, typical for large-scale fMRI data collections, the inflow signal of cardiac-related flow will be aliased and falsely appear as a low frequency modulation, as the Nyquist criteria is not met (23). Developing a robust method to quantify CSF flow from conventional fMRI acquisitions could open new opportunities to investigate the relationship between CSF flow and various cognition- and aging-related metrics that are commonly acquired in ongoing fMRI studies. Such relationships have the potential to provide valuable insights into areas including brain clearance and the progression of neurodegenerative diseases (24).

Here we propose a method that takes spin history into account to quantify CSF flow from the observed inflow effect in fMRI data. The method does not depend on rapid sampling and can be implemented with standard fMRI acquisitions. We validated the method against phantom measurements and applied it to a typical fMRI dataset to quantify the magnitude of cardiac, respiratory and slow vasomotion cycles for CSF flow at the level of foramen magnum, thus directly addressing the question regarding the importance of different physiological cycles to CSF flow.

## Materials and Methods

### Subjects

Sixty-one elderly individuals were randomly selected from the population registry in Umeå Sweden (25). Eight did not complete the fMRI and five were excluded due to poor physiological recordings during the scan (pulse oximeter and respiratory bellows). Thus, the study population consisted of 48 individuals (68-82 years, 19 women), with characteristics summarized in Table 1. Prior to the MRI scans, all participants underwent a medical examination, during which blood pressure was measured in seated position on the left arm. Relevant medical conditions and medications were also noted. Cognition was tested in all participants using a Montreal Cognitive Assessment (MoCA) test.

**Table 1.** Descriptive characteristics of the participants in this study (N = 48). Categorical variables are given as N (%), while continuous variables are presented with mean ± SD.

|  |  |
| --- | --- |
| Age (Years) | 76.1 ± 4.0 |
| Sex (Men/Women) | 29/19 (60/40) |
| BMI (kg/m<sup>2</sup>) | 24.9 ± 3.5 |
| SBP (mmHg) | 135 ± 16 |
| DBP (mmHg) | 75 ± 10 |
| MoCA score, N (%) | 26.2 ± 2.4 |
| Diabetes mellitus, N (%) | 6 (13) |
| Hypertension, N (%) | 28 (58) |
| Hyperlipidemia, N (%) | 25 (52) |
| Atrial fibrillation, N (%) | 10 (21) |
| Transient ischemic attack, N (%) | 3 (6) |

- 3 N, number of participants; SD, standard deviation; BMI, body mass index; SBP, systolic - 4 blood pressure; DBP, diastolic blood pressure; MoCA, Montreal Cognitive Assessment

The study was conducted according to the ethical principles outlined in the Helsinki Declaration, and the Swedish Ethical Review Authority has approved the study (approval: 2020-03710). Informed consent was obtained both written and orally from all participants. To only include participants who could provide informed consent, a minimum MoCA score of 20 out of 30 possible points was required for participation.

### MRI protocol

MRI acquisitions including resting-state fMRI and high-resolution T_1_-weighted volumes were acquired using a 3T scanner (Discovery MR 750; GE Healthcare, Milwaukee, Wisconsin), with a 32-channel head coil. Resting-state fMRI was obtained through a BOLD-contrast sensitive T_2*_-weighted single-shot gradient echo-planar imaging (EPI) sequence. The imaging volumes were collected with 37 axial slices in an interleaved order with a slice thickness of 3.4 mm and 0.5 mm spacing. The imaging parameters were set as follows: TR/TE 2000/30 ms, flip angle 80°, and field of view (FOV) 25 × 25 cm^2^ with in-plane resolution 128 × 128 voxels. The sequence began with 10 dummy scans so that the magnetization had reached a steady state before any data was collected. Furthermore, physiological recordings including pulse and respiration were collected throughout the scanning period (6 min) using a peripheral pulse unit and respiratory bellows.

A baseline anatomical IR-prepped T_1_-weighted volume was collected using a fast spoiled gradient echo (FSPGR) sequence with the following imaging parameters: TI/TR/TE 450/8.2/3.2 ms, flip angle 12°, 1 mm slice thickness, in-plane resolution 0.93 mm, FOV 25 × 25 cm^2^ with 176 axial slices, and parallel imaging factor of 2. Variable flip angle T1-mapping was implemented with FSPGR acquisitions and flip angles 2° and 12° using the same FOV and resolution as the anatomical scan. From these images, subarachnoid space CSF T_1_-times for each participant were estimated (26).

The MRI acquisitions were conducted within a broader quantitative imaging protocol that included intravenous gadolinium injection (25), which was not utilized in the present study. In this protocol the baseline T_1_-weighted volumes were collected before the injection of gadolinium (Dotarem), whereas the resting-state fMRI acquisition was approximately 3 hours after contrast injection. Gadolinium potentially impacts the BOLD signal, but it has been reported that even when acquisition occurs close to the injection time, no statistically significant mean signal change is observed (27). Moreover, while intravenous injections of Gadolinium can have a measurable impact on the T_1_-times of CSF (28) our measurement protocol involved measuring the average T_1_-times before and after the fMRI acquisition. Therefore, the effect of Gadolinium on the T_1_-values used in the model was accounted for.

### fMRI processing

Before any preprocessing, we manually drew regions of interest (ROIs) in the area of foramen magnum on the bottom slice of the fMRI volume to capture the inflow effect of CSF. The manual drawing was based on the difference between each voxel’s maximum and minimum signal intensity, thereby highlighting voxels with significant movement. The ROIs were divided into two classes based on K-means clustering on the EPI signal to separate CSF voxels from non-CSF voxels, where the class containing the most voxel was taken to be CSF voxels. This voxel division was visually inspected to verify a representative segmentation of the CSF region. The signal related to CSF movement was then calculated as a spatial average of all CSF voxels, and the resulting time-series was normalized by division of its mean. For the BOLD signal analysis, realignment, slice timing and co-registration to a T_1_-weighted volume was performed on each fMRI session using SPM12 (29). A whole brain cortical gray matter ROI was created using cortical surface segmentations in FreeSurfer 6.0 (30) and removal of physiological noise was performed using RETROICOR (31). After that, the global BOLD signal in gray matter was extracted from the mean value of all voxels within the gray matter ROI after voxel wise normalization to z-score. The signal was then detrended using a polynomial of degree 2 prior to the application of a band-pass filter (0.01–0.1 Hz), where the processed signal serves to characterize *slow vasomotion* throughout the study. The process of extracting the EPI signal associated with CSF movement from the bottom slice is illustrated in Figure 1. Since high displacements impact the spin history, the framewise displacement was calculated to exclude participants with high movement during the fMRI scan. Here, we excluded participants with a mean framewise displacement greater than 0.5 mm or a maximum framewise displacement greater than 3 mm (N = 5) as motion induced changes in spin history are not considered in the model and may therefore have a negative impact on the fit of the simulated signals.

**Figure 1.**
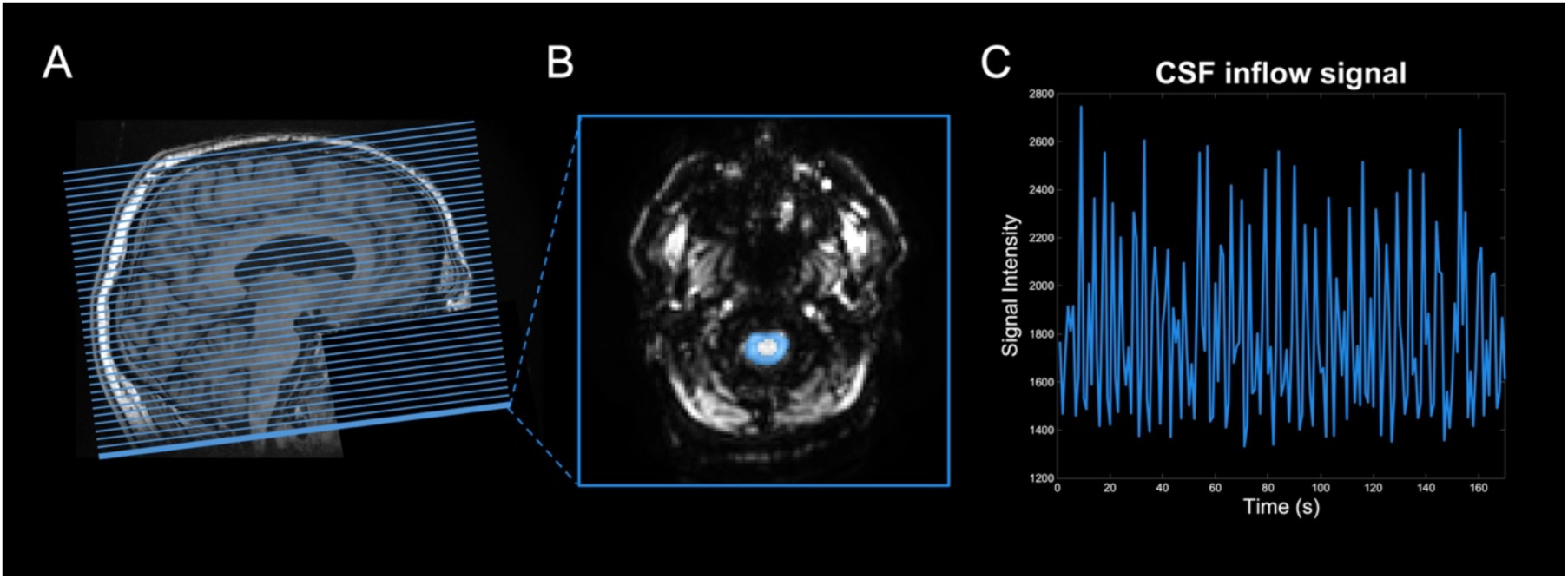
Illustrations of the CSF signal extraction in fMRI. **(A)** T_1_-weighted image showing fMRI slice coverage. **(B)** The bottom fMRI slice, with the CSF region of interest highlighted. This region was used to extract the inflow effect time series shown in **(C)**.

### Modeling CSF dynamics

The CSF dynamics modeling presented in this study is based on the Monro-Kellie doctrine (18), where the total volume of brain, intracranial blood and CSF is assumed to be constant. This suggests that CSF buffers variations in blood volume by flowing in and out of the spinal compartment, which we want to quantify from the EPI signal fluctuations around foramen magnum. The intracranial blood volume is influenced by cardiac pulsations, respiratory-driven pressure changes, and slower fluctuations related to neural activity and autoregulatory mechanisms (vasomotion). Therefore, we constructed a model to mathematically describe the signal fluctuations caused by CSF movement, where the CSF velocity is also assumed to depend on cardiac pulsations, respiratory variations, and slow vasomotion derived from the global BOLD signal. The physiological signals used in the modelling are visualized together in Figure 2. We assumed that the cardiac, respiratory and slow vasomotion components of the CSF velocities are quasi-periodic, captured by the phases calculated from recordings of pulse, respiration and the global BOLD signal.

**Figure 2.**
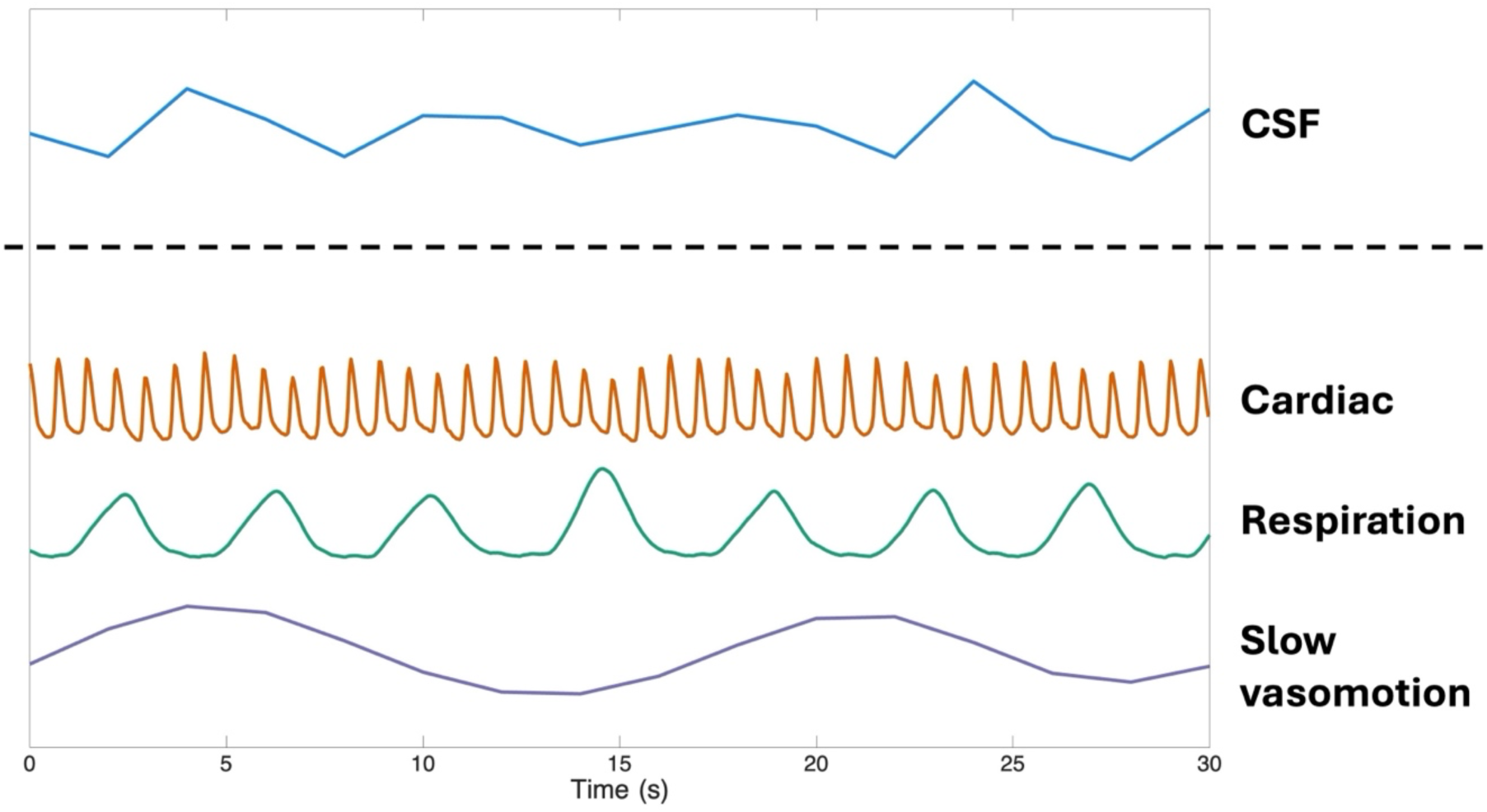
Time-series of physiological signals extracted from the fMRI acquisition. The CSF signal was derived from EPI signal fluctuations within the bottom-slice CSF region of interest (ROI). Cardiac and respiratory signals were recorded via pulse oximetry and respiratory bellows, respectively, while slow vasomotion was estimated from the global BOLD signal. Note that the sampling frequency of the CSF signal fluctuations is that of the fMRI sequence (i.e. 2 s).

The phases are calculated with a previously described method that uses the Hilbert-Transform (HT) of a physiological signal to extract the corresponding phases (32). This approach was selected specifically because CSF oscillations are neither purely sinusoidal nor strictly stationary. The HT approach allows for the extraction of instantaneous phase information despite the natural irregularity and non-sinusoidal morphology of real physiological waves (33). With this method, the cardiac phase (𝜑_𝐶_) was calculated via the HT of the pulse oximetry data. Similarly, the respiratory phase (𝜑_𝑅_) was calculated through the HT of the respiratory bellows data. Lastly, for the slow vasomotion component, we used the time series representing the negative derivative of the global BOLD signal to calculate the phase (𝜑_𝑆𝑉_) via the HT, since this derivative should represent the rate of change in cerebral blood volume (14). Using the assumptions above, the CSF velocity over time (𝑡) was written as,

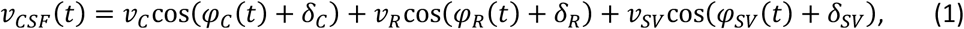

where 𝑣_𝐶_, 𝑣_𝑅_, and 𝑣_𝑆𝑉_ represent the magnitude of the cardiac, respiratory and slow vasomotion components of the CSF velocity. Moreover, 𝛿_𝐶_, 𝛿_𝑅_ and 𝛿_𝑆𝑉_ are phase shifts between the CSF velocity and the cardiac, respiratory and slow vasomotion phases respectively. The CSF displacement was achieved by integrating 𝑣_𝐶𝑆𝐹_. A least squares estimation of the velocities and phase shifts for each velocity component was acquired by,

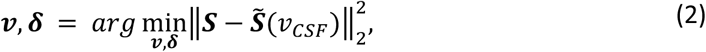

where 𝑺 is the measured inflow signal, 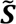 is the simulated signal, 𝒗 = {𝑣_𝐶_, 𝑣_𝑅_, 𝑣_𝑆𝑉_}, and 𝜹 = {𝛿_𝐶_, 𝛿_𝑅_, 𝛿_𝑆𝑉_}. The optimization problem was minimized using particle swarm (34) with upper and lower bounds on the velocities corresponding to ±50 mm/s for each component.

### Simulations of CSF inflow signal

An illustration of the simulation is provided in Figure 3, as well as in the supplementary movie. The simulations began with setting *N* equispaced grid points (i.e. the oscillating ensemble) in a region around the bottom slice of the imaging volume. Each of the grid points were assigned an initial magnetization 𝑀_0_ along the main magnetic field, representing elements of CSF. After that, the position of each element was updated according to 𝑣_𝐶𝑆𝐹_(𝑡), where a plug flow was used as flow profile of the CSF flow. The inflow effect was simulated by continuously updating the magnetization of moving CSF throughout the fMRI sequence, based on solutions to the Bloch equations (35) and by considering excitation of multiple slices. With the main magnetic field along the z-direction, the initial unperturbed net magnetization is along the z-direction with magnitude 𝑀_0_. Directly after the application of a radiofrequency (RF) pulse of flip angle 𝜃, the magnetization parallel to the main magnetic field (𝑀_𝑧_) and the transverse magnetization (𝑀_⊥_) become

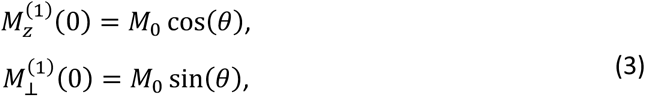

where the superscript indicates the number of RF pulses applied. From the Bloch equations, the relaxation of the magnetizations after RF pulses become

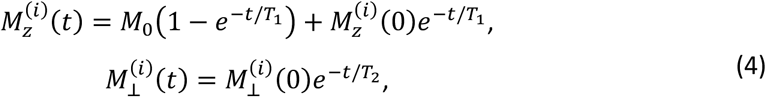

where 𝑇_1_ and 𝑇_2_ are the T_1_ and T_2_ relaxation times. A general expression for the parallel and transverse magnetization after *i* RF pulses is

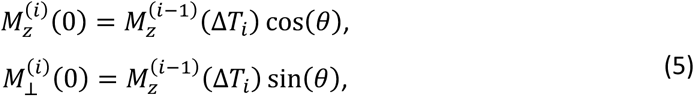

where Δ𝑇_𝑖_ is the time between the two succeeding pulses experienced by an element and will be equal to TR only if both excitations occur within the same imaging slice. However, due to CSF motion, the element may undergo excitations in other slices, resulting in Δ𝑇_𝑖_ differing from TR.

**Figure 3.**
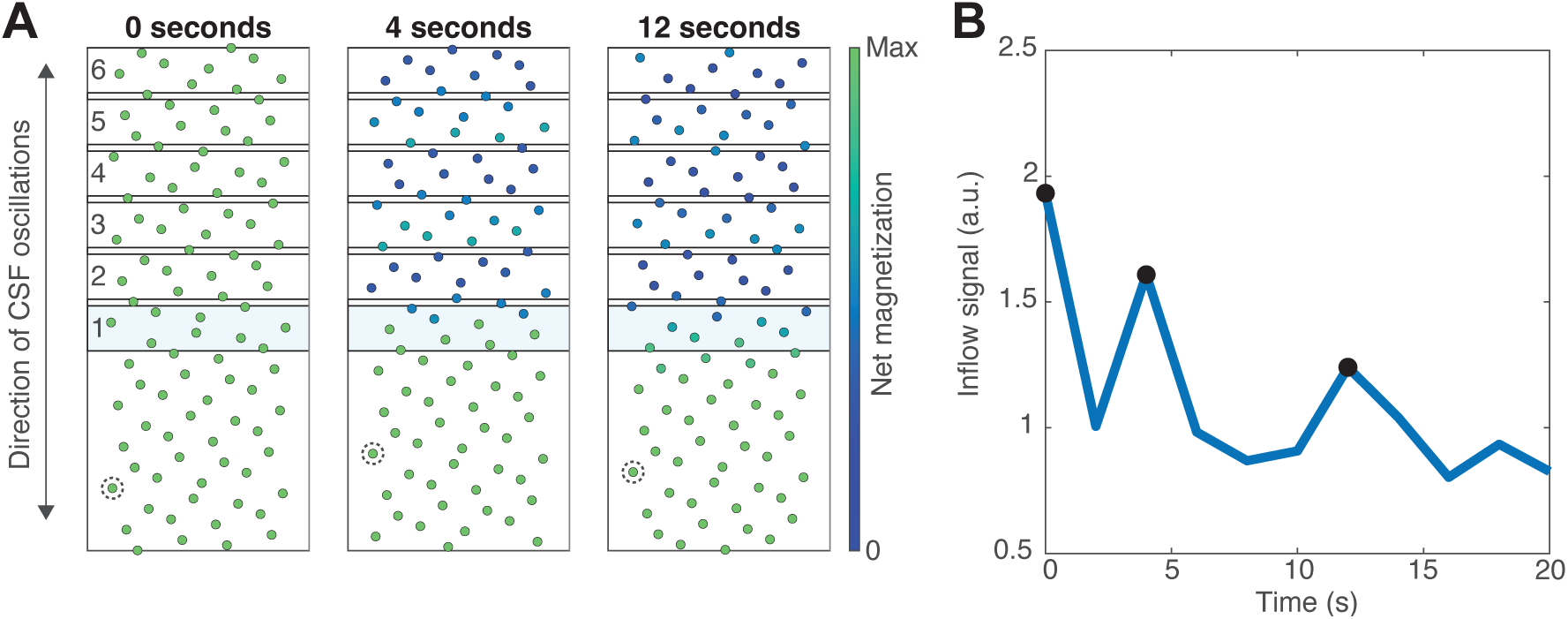
Visualization of the simulation approach. (A) Magnetizations of particles in an ensemble that oscillates in a direction perpendicularly to the imaging slices. Magnetization is provided at three example time points where the edge slice is being excited (illustrated by the light blue color). To help visualize the motion, one particle has been highlighted by a dashed line. (B) The signal in the edge slice, with the example time points in (A) marked.

The magnetizations of the elements were updated according to the Equations 3-5 together with the settings of the fMRI sequence, which includes the excitation of multiple slices. Physiologically, within the CSF space, there is some degree of dispersion that would cause blurring of the magnetization between adjacent positions. Therefore, the parallel magnetization at each time step and position was convolved with a Gaussian kernel corresponding to such a process with a dispersion coefficient of 6 cm^2^/min. This was chosen based on the phantom validation and is in the same order of magnitude as previously reported dispersion coefficient of spinal CSF (36). The T_1_-value for CSF was set to the cohort average 4738 ± 186 ms, while the T_2_-value was set 1600 ms based on the literature (37). The signal at the bottom slice was acquired by the mean transverse magnetization of all elements localized within the slice, evaluated at TE for each excitation of the bottom slice. Moreover, the simulated signal was normalized by its mean, eliminating the need to determine 𝑀_0_.

### Model validation

The effects of overfitting on the model were examined by fitting the model when interchanging each participant’s cardiac, respiratory and slow vasomotion recordings by another participant’s recordings. For each participant, the physiological recordings were interchanged with a participant that displayed a low correlation of the physiological recordings, using a metric that was calculated as the maximum absolute cross-correlation between each of the three physiological signals. However, since we want the interchange of gatings to be a permutation, the gatings swaps were therefore performed using Munkres algorithm (38). The explained variance (R^2^) from model fits using swapped recordings were presented with those derived from the true recordings to compare the model’s ability to describe the EPI-inflow signal when using both true and false gating data. Furthermore, given that the model depends on the selection of the T_1_ relaxation time, we examined the effect of this parameter by comparing CSF stroke volumes obtained when the group average T_1_-time (4738 ms) was used compared to the individually calculated T_1_-times.

The model was further validated using phantoms mimicking oscillatory flow of CSF. Here a hollow cylinder with a length of 9.2 cm and inner/outer diameter of 2.0/4.3 cm was cast in agar (30 g/L) to avoid water/plastic interfaces. An in house developed syringe pump was used to generate oscillatory flow inside the hollow cylinder, which followed a sine-curve with a fixed oscillatory frequency. The frequency of the oscillatory flow was set to 0.93 Hz, 0.23 Hz, and 0.077 Hz respectively to mimic cardiac, respiratory, and slow vasomotion cycles. For the phantom measurements, we used water as flowing medium, and the imaging parameters of the fMRI acquisitions were the same as the in vivo scan except for the scanning time that was now decreased to 2 min. The evaluation was performed with 6 different velocity amplitudes for each of the three frequencies, with cardiac velocities ranging between 0 and 20 mm/s, respiratory within 0 and 5 mm/s, and slow vasomotion within 0 and 2.5 mm/s. The simulations of the inflow signal for the oscillating flow in the phantom were repeatedly run with different dispersion coefficients to evaluate the robustness of the model.

The assumption of that a plug flow is representative to the CSF flow profile was validated by simulating signal variations with different flow profiles and fitting a plug flow to the corresponding signal change. Here we simulated signals corresponding to flow profiles of a laminar flow inside a cylinder, and a laminar flow inside an annulus with outer radius 3 times larger than the inner radius, and a plug flow. The simulations were repeatedly run with mean velocities ranging from 1 mm/s to 25 mm/s, and the fitted plug velocities were compared to the mean velocities of the different flow profiles.

### Statistics

All statistical analyses were conducted in MATLAB (version 9.13.0. Natick, Massachusetts: The MathWorks Inc.). Normality of the reported variables was assessed using skewness and kurtosis by calculating z-scores, with values within ±1.96 indicating normality. Normal variables are reported as mean ± standard deviation, while non-normal variables are reported with median and interquartile range. Pearson correlation coefficient was used to measure associations between two variables and p-values were considered significant at the 0.05 level.

## Results

### CSF flow rates and volumes

When applied to a whole-brain fMRI data set of N=48 individuals (68-82 years, 19 women), quantitative estimates of the CSF flow at the cranio-cervical junction could be obtained in all subjects. The simulated and measured CSF inflow displayed R^2^ values of 0.57 [0.47, 0.67] supporting the appropriateness of the model, see Figure 4A for the distribution of individual R^2^ values. Figure 4A also contains the outcome of a negative-control test, where the inflow signal was deliberately matched to cardiac, respiratory and BOLD signals of another subject, verifying that a good fit could not be obtained by chance. A typical fit is displayed in Figure 4B (R^2^ = 0.64). The estimated velocity amplitudes of the cohort were 12.0 [8.1, 18.4] mm/s for the cardiac component, 1.21 [0.76, 2.14] mm/s for the respiratory component, and 0.18 [0.11, 0.24] mm/s for the slow vasomotion component (Figure 4C). After factoring in the area of the CSF region, the corresponding flow rates and stroke volumes were calculated to be 3.03 [2.43, 4.63] mL/s and 0.77 [0.57, 1.09] mL for the cardiac cycle, 0.32 [0.21, 0.61] mL/s and 0.38 [0.26, 0.88] mL for the respiratory cycle, and 0.049 [0.024, 0.070] mL/s and 0.26 [0.14, 0.39] mL for the slow vasomotion cycle (Figure 4D and 4E). Moreover, the agreements between the stroke volumes estimated using a fixed T_1_-time versus the individually set T_1_-times were high, as indicated by small systematic differences (1%, 3% and 1% for cardiac, respiratory, and slow vasomotion, respectively), and high correlations values (0.987 0.996 and 0.971 for cardiac, respiratory, and slow vasomotion, respectively) (see Figure 5).

**Figure 4.**
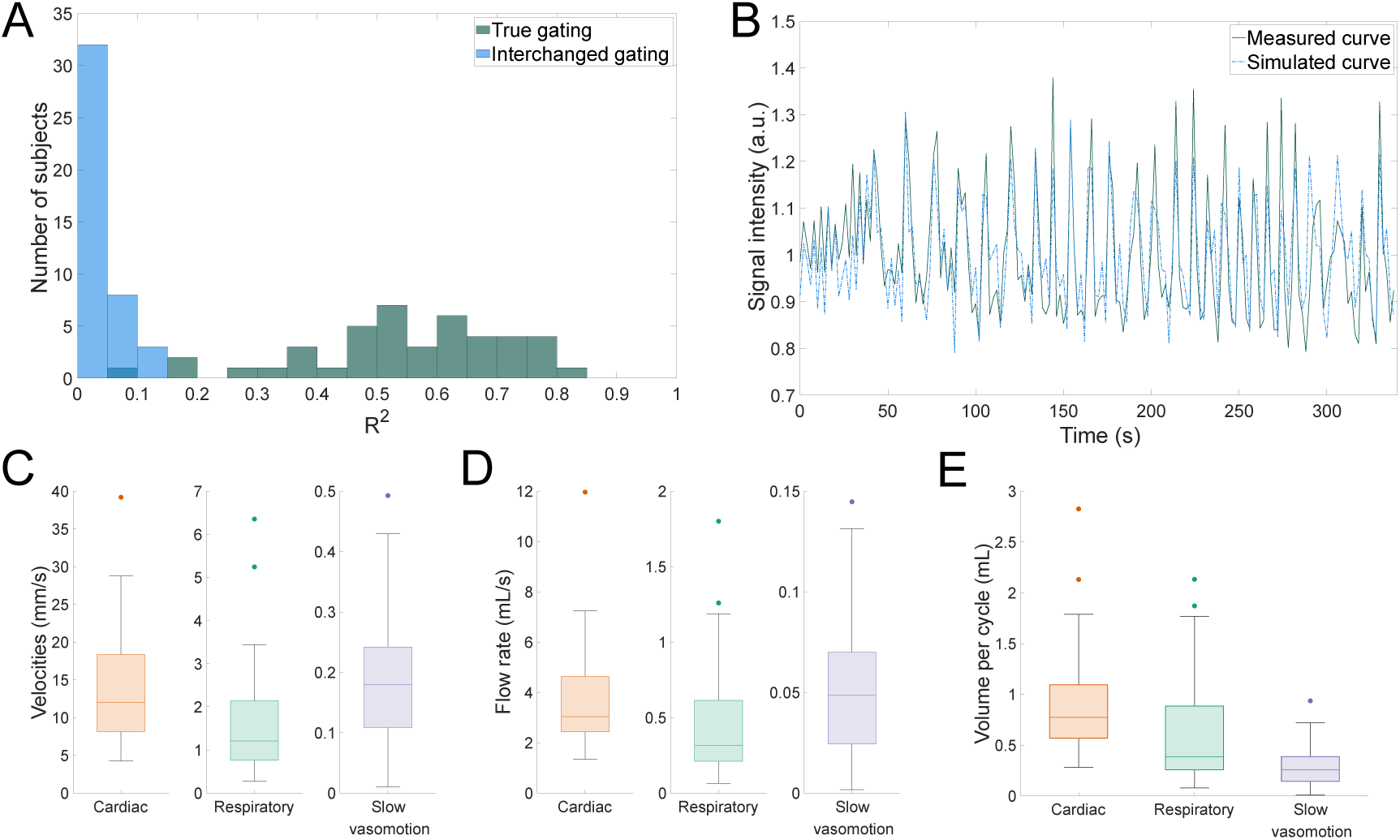
Results from the simulations run on the participants of this study (N = 43). (A) Correlation coefficients between simulated and measured CSF signal for each subject, both with each participants true gating and interchanged gatings. (B) Example fit of the CSF signal. (C) Estimated velocities for each subject and physiological component. Note the different scales on the vertical axes. (D) Estimated flow rates for each subject and physiological component. Note the different scales on the vertical axes. (E) Estimated stroke volumes over each of the physiological cycles for each subject.

**Figure 5.**
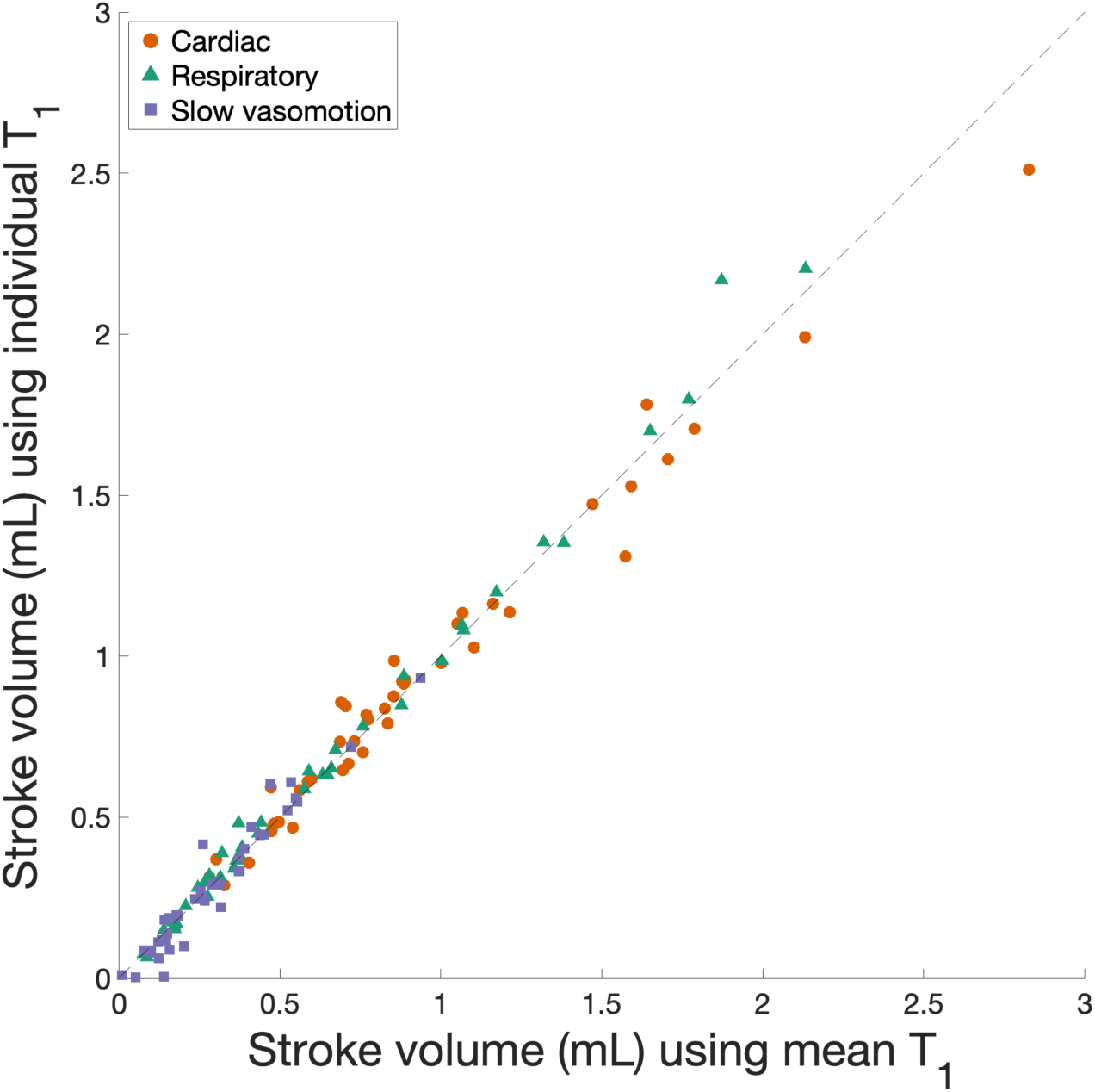
Comparison of estimated CSF stroke volumes when using the cohort mean T_1_-time versus individual participant T_1_-time in the modeling. The figure presents the results across the three physiological domains.

### Phantom validation

The correlation coefficients and linear fits between estimated velocity and the applied pump velocity for all simulations are summarized in Table 2, with all the correlations being statistically significant. Because the slow vasomotion frequency exhibited a strong linear relationship with velocities up to 1.5 mm/s, but weakened at higher velocities, we included a separate column in Table 2 presenting data limited to this velocity range. The estimated velocities for each of the three frequencies together with the applied pump velocity are displayed in Figure 6A-C for the simulations with a dispersion coefficient of 6.00 cm^2^/min.

**Figure 6.**
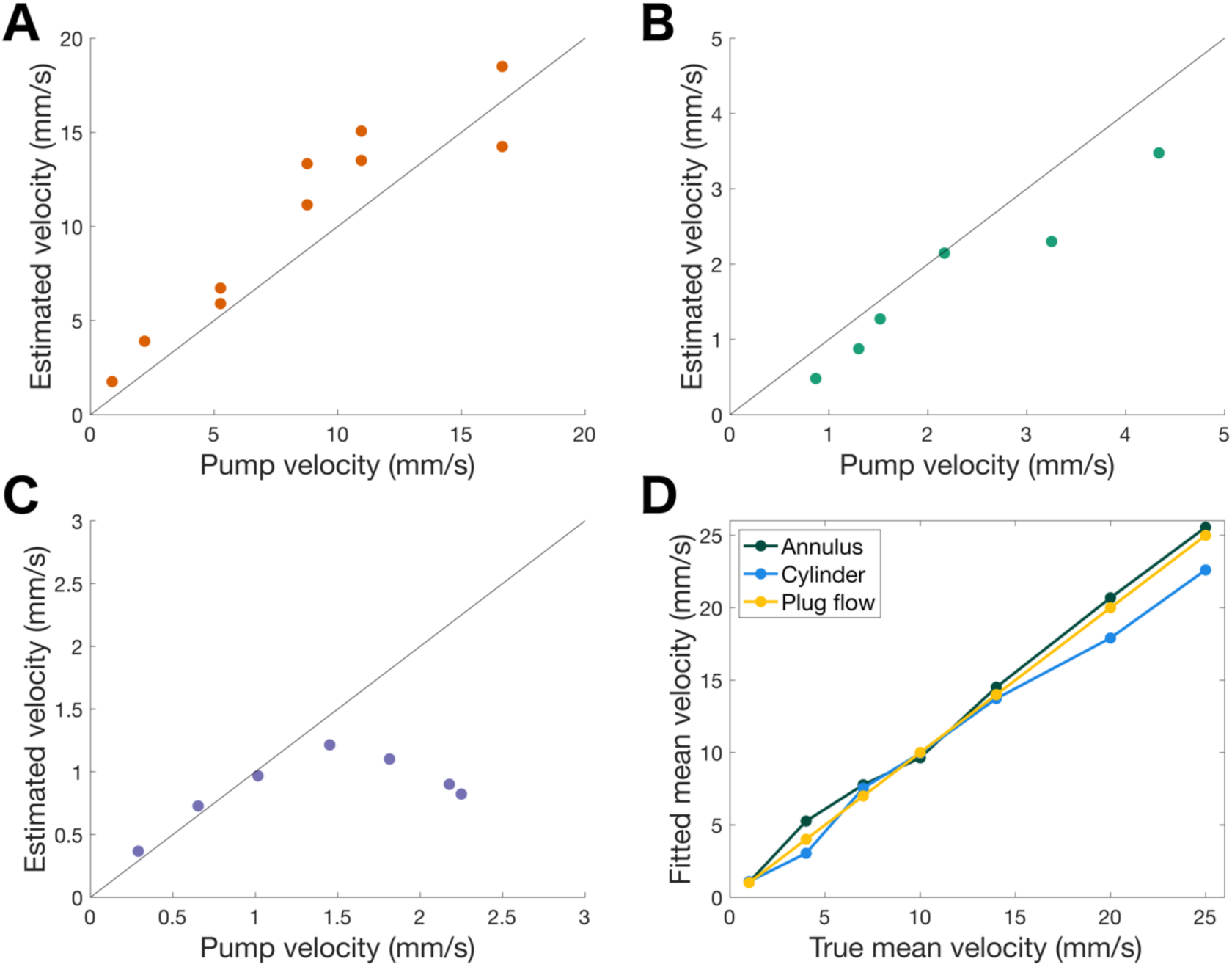
Results from phantom measurements when using a dispersion coefficient of 6.00 cm^2^/min. (A) Estimated velocities vs pump velocities for cardiac reflected frequency (0.93 Hz). (B) Estimated velocities vs pump velocities for respiratory reflected frequency (0.23 Hz). (C) Estimated velocities vs pump velocities for slow vasomotion reflected frequency (0.077 Hz). (D) Mean velocities obtained by fitting a plug flow to simulated signals of different flow profiles plotted against the underlying mean velocities.

**Table 2.** Results from phantom measurements when using different values of the dispersion coefficient. The correlation coefficient between estimated and measured fMRI signal (r) and the linear fit with intercept α and slope β are presented for each dispersion coefficient and each operating frequency.

| | Cardiac (0.93 Hz) | | | Respiratory (0.23 Hz) | | | Slow vasomotion (0.077 Hz) | | | Slow vasomotion (0.077 Hz)<br>(Velocities $\leq 1.5$ mm/s) | | |
| --- | --- | --- | --- | --- | --- | --- | --- | --- | --- | --- | --- | --- |
| Dispersion<br>coefficient<br>(cm<sup>2</sup>/min) | $r$ | $\alpha$ | $\beta$ | $r$ | $\alpha$ | $\beta$ | $r$ | $\alpha$ | $\beta$ | $r$ | $\alpha$ | $\beta$ |
| 2.00 | 0.9270 | 1.7316 | 0.6500 | 0.9641 | 0.0148 | 0.5592 | 0.5735 | 0.4308 | 0.1530 | 0.9899 | 0.1547 | 0.5279 |
| 3.00 | 0.9277 | 2.0623 | 0.7445 | 0.9729 | -0.0573 | 0.6689 | 0.5685 | 0.4820 | 0.1742 | 0.9835 | 0.1698 | 0.5966 |
| 4.00 | 0.9281 | 2.1183 | 0.8338 | 0.9670 | -0.0097 | 0.6959 | 0.5651 | 0.5265 | 0.1842 | 0.9862 | 0.1876 | 0.6446 |
| 5.00 | 0.9214 | 2.3760 | 0.8704 | 0.9682 | -0.0493 | 0.7724 | 0.5808 | 0.5423 | 0.2072 | 0.9882 | 0.1812 | 0.6954 |
| 6.00 | 0.9368 | 2.2039 | 0.9504 | 0.9724 | -0.0559 | 0.8100 | 0.5767 | 0.5816 | 0.2107 | 0.9907 | 0.2061 | 0.7197 |
| 7.00 | 0.9303 | 2.5455 | 1.0044 | 0.9658 | -0.0865 | 0.8539 | 0.5716 | 0.6094 | 0.2193 | 0.9904 | 0.2117 | 0.7590 |
| 8.00 | 0.9207 | 2.7232 | 1.0339 | 0.9598 | -0.0124 | 0.8410 | 0.5854 | 0.6179 | 0.2279 | 0.9828 | 0.2273 | 0.7563 |

The estimated mean velocities obtained by fitting a plug flow to the signals of different flow profiles showed strong agreement with the corresponding mean velocities of the underlying flow profiles. The correlation coefficients between the fitted and actual mean velocities were: 0.996 for laminar flow in a cylinder, 0.998 for laminar flow in an annulus. Per design, there was a perfect correlation with plug flow. A summary of these comparisons is presented in Figure 6D.

## Discussion and Conclusions

We propose a method to characterize cardiac, respiratory and slow vasomotion velocity components of CSF motion compatible with a standard whole-brain fMRI implementation, where CSF is captured in the bottom slice of the imaging volume. Using this method, we could show that CSF flow at the foramen magnum was dominated by cardiac related oscillations, with lower stroke-volumes for the respiratory and slow vasomotion cycles. The method was inspired by recent advancements in utilizing the inflow-related signal enhancements that develop because of CSF oscillations in and out of the imaging volume (e.g. at the fourth ventricle or at the foramen magnum) (14–16). However, the approach presented here provides some major benefits compared to previous studies. Firstly, the method estimated *quantitative* values of CSF velocities, based on a model that does not depend on specific imaging parameters and is verified with phantom experiments. Although qualitative assessments are useful for identifying population trends, they lack the quantitative precision required for individual comparisons. Since flow is a product of velocity and area, two subjects with similar signal intensities can have vastly different flow rates due to variations in foramen magnum geometry. Furthermore, our results demonstrated that the method is robust using a typical rather than measured T_1_-time of CSF, supporting use in datasets where the CSF T_1_-acqusition is not available. Secondly, using recordings of cardiac and respiration cycles, individual contributions from physiological cycles could be estimated. Since typical fMRI sampling frequency does not meet the Nyquist criterion, such decompositions cannot be made from the raw signal itself. The method presented in the current study is intended to offer means of retrospectively assessing CSF flow in large-scale studies that included fMRI acquisitions where edge-slice CSF is visible, as would be expected in studies employing whole-brain fMRI. If physiological recordings are missing or of insufficient quality, deep-learning frameworks (39) can potentially be used provide robust estimates of these signals directly from the fMRI data.

In the present study we applied the method to analyze CSF flow and stroke volumes at the cranio-cervical junction, where we found median cardiac velocity amplitudes of 12.0 mm/s and respiratory velocity amplitudes of 1.21 mm/s. Previous literature has reported CSF velocities and stroke volumes in this region using PC-MRI; with cardiac velocities by Yildiz et al. 2017 and 2022 to be 9.7 ± 3.3 mm/s (40) and 17.5 ± 4.0 mm/s (41) respectively, compared to Takizawa et al. that reported cardiac velocities of ∼10 ± 5 mm/s (42). The corresponding respiratory velocities in the same studies were 5.8 ± 4.0 mm/s (40), 6.8 ± 3.4 mm/s (41) and ∼3 ± 1 mm/s (42). However, in the study by Yildiz et al. 2022 they consider a single voxel approach that captures peak velocity, rather than average CSF velocities. Wåhlin et al. have reported cardiac stroke volumes at cervical level to be 0.773 ± 0.231 mL (1) while Liu et al. presented values of ∼0.6 mL (43), both correspond very well with our estimate of 0.77 mL. The respiratory stroke volumes at cervical level during guided breathing have been reported to be 0.99 mL in mean and range from 0.54 mL to 1.66 mL (44), which are higher than our estimations obtained during free breathing (0.38 mL). However, the breathing *mode* may affect the CSF modulation (45), where guided breathing likely increases the effects from respiration compared to free breathing. Importantly, these effects may vary with age (46), and since the participants in our cohort were older than those in the studies used for comparison, some caution in the comparisons of abolute values is warranted.

For the slow vasomotion component of the CSF velocity at foramen magnum, no established reference values exist. However, a crude estimation of CSF velocity can be derived from intracranial pressure (ICP) recordings when analyzing slow waves variations of ICP. The velocity due to oscillatory ICP changes is given by 𝑣 = 2𝜋𝑓 · Δ𝑃 · 𝐶 · 𝑟_𝑐_ / 𝐴, where 𝑓 is the operating frequency of the ICP waves, Δ𝑃 is the amplitude of the pressure changes, 𝐶 denotes the intracranial compliance, 𝑟_𝑐_ is the spinal fraction of the compliance, and 𝐴 is the area of the CSF flow pathway at foramen magnum. For this calculation we used 𝑓 = 0.05 Hz, Δ𝑃 = 0.8 mmHg (47), 𝐶 = 1.15 mL/mmHg (48), 𝑟_𝑐_ = 35% (4), and 𝐴 = 3.6 cm^2^ (49). By combining these reference values, the estimated slow vasomotion CSF velocity is approximately 0.28 mm/s. This velocity is of the same order of magnitude as observed in the present study. Moreover, the upper limit of the strong linear relationship observed for the slow vasomotion component in the phantom measurements is approximately five times the estimated velocity, indicating that our method performs reliably within the expected physiological range.

Previous studies have presented an approach to utilize the inflow effect to relate CSF flow and brain activity by calculating the correlation between the CSF signal and global BOLD signal. This method has been successful in linking a reduced coupling between CSF flow and brain activity to neurodegenerative diseases and cognitive decline (15,16). Nevertheless, these results may not fully capture the magnitude or change of the underlying CSF flow, as the correlation measure only reflects degree of temporal coupling. A quantitative approach to estimate CSF flow from the inflow signal has been proposed where the signal equations derived by Gao et al. (50) have been used to translate the signal fluctuations to CSF flow (51). However, a limitation with their approach is that it does not capture the oscillating behavior of CSF flow as the signal equations are derived for unidirectional flow and only considers single-slice excitation. Another approach is to use PC-MRI and BOLD fMRI in an interleaved fashion (52), which can capture both flow measurements and the BOLD signal accurately. Such specialized imaging may however not always be feasible (e.g. in ongoing longitudinal aging and dementia studies), whereby a method to retain similar information from standard imaging is desirable. Moreover, a recently presented approach based on multiple slice excitation together with artificial intelligence has shown promising results to translate the inflow signal to velocities when using fMRI data with short repetition times (53).

### Limitations

As with all dynamic MRI imaging, the proposed approach is sensitive to motion of the subjects since it induces spin-history changes that are difficult to correct for (54). Because of this, we have excluded participants with high movement during the scanning since they are expected to yield unreliable results for this kind of analysis. The fMRI acquisition is also sensitive to B_0_ fluctuations which may mainly be affected by respiration. However, the respiratory B_0_ fluctuations at cervical level 1 and the brain have been shown to be relatively small (55). Moreover, the model assumes the CSF velocity distribution to be well approximated by a plug flow, although the underlying velocity distribution is unknown. However, as simulations showed that the fitted velocities of plug flow display a high correlation to the underlying mean velocities for various flow profiles, the impact of this shortcoming is probably limited. Furthermore, given that this method is intended for the retrospective analysis of fMRI cohorts, the bottom slice may not always be perfectly orthogonal to the foramen magnum. However, based on the principle of mass conservation, any slice obliquity is expected to increase the cross-sectional area, effectively compensating for the decrease in the estimated velocity component. Consequently, the calculated volumetric flow rate theoretically remains unaffected in such cases. Additionally, Equation 1 does not include a term for the net flow of CSF. Previous research suggests that the CSF production rate in humans is approximately 0.5 mL/min (56). If 50% of the produced CSF is assumed to pass through foramen magnum (57), with an area of 3.6 cm² (49), the resulting net velocity would be about 0.01 mm/s. Since this net velocity is substantially lower than the oscillatory velocities, it was considered negligible for the model and therefore disregarded in this study to keep the model complexity down. Lastly, the model was only validated using one set of imaging parameters representative of typical large-scale fMRI-datasets. However, we see no intrinsic limitation in the approach that would render the method infeasible for variations in parameter settings (e.g. TR, slice-thickness, slice order, between-slice acceleration).

## Conclusion

By modeling spin-history we were able to quantify CSF flow from the inflow effect in fMRI data. The cardiac, respiratory and slow vasomotion flow rates could be separated and cyclic volume variations (i.e. stroke volume) associated with each cycle could be calculated. CSF motion was dominated by the cardiac cycle since this cycle was faster and of greater stroke-volume compared to the other two cycles. Importantly, the proposed framework requires no specialized pulse sequence. It was designed to be flexible and readily adaptable to standard fMRI acquisitions, provided that physiological signals are available along with edge-slice CSF coverage.

## Acknowledgements

This study was supported by grants from the Swedish Research Council (grant number 2022-04263), the Swedish Heart-Lung Foundation (grant number 20210653), and from the Swedish Foundation for Strategic Research (grant number RMX18-0152). Gemini (Google LLC, Mountain View, CA, USA) was used to assist with sentence structuring during manuscript preparation. The authors take full responsibility for all text in the manuscript.

## Data Availability Statement

The data underlying this article will be shared on reasonable request to the corresponding author.

## References

1. Wåhlin A, Ambarki K, Hauksson J, Birgander R, Malm J, Eklund A. Phase contrast MRI quantification of pulsatile volumes of brain arteries, veins, and cerebrospinal fluids compartments: repeatability and physiological interactions. Journal of Magnetic Resonance Imaging 2012;35(5):1055–1062.

2. Vijayakrishnan Nair V, Kish BR, Inglis B, Yang H-C, Wright AM, Wu Y-C, Zhou X, Schwichtenberg AJ, Tong Y. Human CSF movement influenced by vascular low frequency oscillations and respiration. Front Physiol 2022;13:940140.

3. Kazimierska A, Kasprowicz M, Czosnyka M, Placek MM, Baledent O, Smielewski P, Czosnyka Z. Compliance of the cerebrospinal space: comparison of three methods. Acta Neurochir (Wien) 2021;163:1979–1989.

4. Wåhlin A, Ambarki K, Birgander R, Alperin N, Malm J, Eklund A. Assessment of craniospinal pressure-volume indices. American journal of neuroradiology 2010;31(9):1645–1650.

5. Rasmussen MK, Mestre H, Nedergaard M. Fluid transport in the brain. Physiological Reviews 2022;102(2):1025–1151.

6. Iliff JJ, Wang M, Liao Y, Plogg BA, Peng W, Gundersen GA, Benveniste H, Vates GE, Deane R, Goldman SA. A paravascular pathway facilitates CSF flow through the brain parenchyma and the clearance of interstitial solutes, including amyloid β. Science translational medicine 2012;4(147):147ra111–147ra111.

7. Jessen NA, Munk ASF, Lundgaard I, Nedergaard M. The glymphatic system: a beginner’s guide. Neurochem Res 2015;40:2583–2599.

8. Mestre H, Tithof J, Du T, Song W, Peng W, Sweeney AM, Olveda G, Thomas JH, Nedergaard M, Kelley DH. Flow of cerebrospinal fluid is driven by arterial pulsations and is reduced in hypertension. Nature communications 2018;9(1):4878.

9. Iliff JJ, Wang M, Zeppenfeld DM, Venkataraman A, Plog BA, Liao Y, Deane R, Nedergaard M. Cerebral arterial pulsation drives paravascular CSF–interstitial fluid exchange in the murine brain. Journal of Neuroscience 2013;33(46):18190–18199.

10. Ozturk B, Koundal S, Al Bizri E, Chen X, Gursky Z, Dai F, Lim A, Heerdt P, Kipnis J, Tannenbaum A. Continuous positive airway pressure increases CSF flow and glymphatic transport. JCI insight 2023;8(12).

11. Dreha-Kulaczewski S, Joseph AA, Merboldt K-D, Ludwig H-C, Gärtner J, Frahm J. Inspiration is the major regulator of human CSF flow. Journal of neuroscience 2015;35(6):2485–2491.

12. van Veluw SJ, Hou SS, Calvo-Rodriguez M, Arbel-Ornath M, Snyder AC, Frosch MP, Greenberg SM, Bacskai BJ. Vasomotion as a driving force for paravascular clearance in the awake mouse brain. Neuron 2020;105(3):549–561. e545.

13. Jiang-Xie L-F, Drieu A, Bhasiin K, Quintero D, Smirnov I, Kipnis J. Neuronal dynamics direct cerebrospinal fluid perfusion and brain clearance. Nature 2024:1–8.

14. Fultz NE, Bonmassar G, Setsompop K, Stickgold RA, Rosen BR, Polimeni JR, Lewis LD. Coupled electrophysiological, hemodynamic, and cerebrospinal fluid oscillations in human sleep. Science 2019;366(6465):628–631.

15. Han F, Chen J, Belkin-Rosen A, Gu Y, Luo L, Buxton OM, Liu X, Initiative AsDN. Reduced coupling between cerebrospinal fluid flow and global brain activity is linked to Alzheimer disease–related pathology. PLoS Biol 2021;19(6):e3001233.

16. Han F, Brown GL, Zhu Y, Belkin-Rosen AE, Lewis MM, Du G, Gu Y, Eslinger PJ, Mailman RB, Huang X. Decoupling of global brain activity and cerebrospinal fluid flow in Parkinson’s disease cognitive decline. Mov Disord 2021;36(9):2066–2076.

17. Kiviniemi V, Wang X, Korhonen V, Keinänen T, Tuovinen T, Autio J, LeVan P, Keilholz S, Zang Y-F, Hennig J. Ultra-fast magnetic resonance encephalography of physiological brain activity–glymphatic pulsation mechanisms? Journal of Cerebral Blood Flow & Metabolism 2016;36(6):1033–1045.

18. Mokri B. The Monro–Kellie hypothesis: applications in CSF volume depletion. Neurology 2001;56(12):1746–1748.

19. Sudlow C, Gallacher J, Allen N, Beral V, Burton P, Danesh J, Downey P, Elliott P, Green J, Landray M. UK biobank: an open access resource for identifying the causes of a wide range of complex diseases of middle and old age. PLoS Med 2015;12(3):e1001779.

20. Barch DM, Burgess GC, Harms MP, Petersen SE, Schlaggar BL, Corbetta M, Glasser MF, Curtiss S, Dixit S, Feldt C. Function in the human connectome: task-fMRI and individual differences in behavior. Neuroimage 2013;80:169–189.

21. Casey BJ, Cannonier T, Conley MI, Cohen AO, Barch DM, Heitzeg MM, Soules ME, Teslovich T, Dellarco DV, Garavan H. The adolescent brain cognitive development (ABCD) study: imaging acquisition across 21 sites. Dev Cogn Neurosci 2018;32:43–54.

22. Gao J-H, Liu H-L. Inflow effects on functional MRI. Neuroimage 2012;62(2):1035–1039.

23. Huotari N, Raitamaa L, Helakari H, Kananen J, Raatikainen V, Rasila A, Tuovinen T, Kantola J, Borchardt V, Kiviniemi VJ. Sampling rate effects on resting state fMRI metrics. Front Neurosci 2019;13:279.

24. Jiang-Xie L-F, Drieu A, Kipnis J. Waste clearance shapes aging brain health. Neuron 2025;113(1):71–81.

25. Vikner T, Garpebring A, Björnfot C, Nyberg L, Malm J, Eklund A, Wåhlin A. Blood–brain barrier integrity is linked to cognitive function, but not to cerebral arterial pulsatility, among elderly. Scientific Reports 2024;14(1):15338.

26. Fram EK, Herfkens RJ, Johnson GA, Glover GH, Karis JP, Shimakawa A, Perkins TG, Pelc NJ. Rapid calculation of T1 using variable flip angle gradient refocused imaging. Magn Reson Imaging 1987;5(3):201–208.

27. Naganawa S, Nihashi T, Fukatsu H, Ishigaki T, Aoki I. Pre-surgical mapping of primary motor cortex by functional MRI at 3 T: effects of intravenous administration of Gd-DTPA. Eur Radiol 2004;14:112–114.

28. Wåhlin A, Garpebring A, Mogensen K, Vikner T, Björnfot C, Qvarlander S, Malm J, Eklund A. Quantitative imaging of brain to cerebrospinal fluid molecular clearance. 2022.

29. Penny WD, Friston KJ, Ashburner JT, Kiebel SJ, Nichols TE. Statistical parametric mapping: the analysis of functional brain images: Elsevier; 2011.

30. Fischl B. FreeSurfer. Neuroimage 2012;62(2):774–781.

31. Glover GH, Li TQ, Ress D. Image-based method for retrospective correction of physiological motion effects in fMRI: RETROICOR. Magnetic Resonance in Medicine: An Official Journal of the International Society for Magnetic Resonance in Medicine 2000;44(1):162–167.

32. Söderström P, Eklund A, Karalija N, Andersson BM, Riklund K, Bäckman L, Malm J, Wåhlin A. Respiratory influence on cerebral blood flow and blood volume–A 4D flow MRI study. Journal of Cerebral Blood Flow & Metabolism 2025;45(8):1531–1542.

33. Kralemann B, Frühwirth M, Pikovsky A, Rosenblum M, Kenner T, Schaefer J, Moser M. In vivo cardiac phase response curve elucidates human respiratory heart rate variability. Nature communications 2013;4(1):2418.

34. Bonyadi MR, Michalewicz Z. Particle swarm optimization for single objective continuous space problems: a review. Evol Comput 2017;25(1):1–54.

35. Bloch F. Nuclear induction. Physical review 1946;70(7-8):460.

36. Ayansiji AO, Gehrke DS, Baralle B, Nozain A, Singh MR, Linninger AA. Determination of spinal tracer dispersion after intrathecal injection in a deformable CNS model. Front Physiol 2023;14:1244016.

37. Drake-Pérez M, Delattre B, Boto J, Fitsiori A, Lovblad K-O, Boudabbous S, Vargas M. Normal values of magnetic relaxation parameters of spine components with the synthetic MRI sequence. American Journal of Neuroradiology 2018;39(4):788–795.

38. Munkres J. Algorithms for the assignment and transportation problems. Journal of the society for industrial and applied mathematics 1957;5(1):32–38.

39. Bayrak RG, Hansen CB, Salas JA, Ahmed N, Lyu I, Mather M, Huo Y, Chang C. DeepPhysioRecon: Tracing peripheral physiology in low frequency fMRI dynamics. Imaging Neuroscience 2025;3:IMAG. a. 163.

40. Yildiz S, Thyagaraj S, Jin N, Zhong X, Heidari Pahlavian S, Martin BA, Loth F, Oshinski J, Sabra KG. Quantifying the influence of respiration and cardiac pulsations on cerebrospinal fluid dynamics using real-time phase-contrast MRI. Journal of Magnetic Resonance Imaging 2017;46(2):431–439.

41. Yildiz S, Grinstead J, Hildebrand A, Oshinski J, Rooney WD, Lim MM, Oken B. Immediate impact of yogic breathing on pulsatile cerebrospinal fluid dynamics. Scientific reports 2022;12(1):10894.

42. Takizawa K, Matsumae M, Sunohara S, Yatsushiro S, Kuroda K. Characterization of cardiac-and respiratory-driven cerebrospinal fluid motion based on asynchronous phase-contrast magnetic resonance imaging in volunteers. Fluids and Barriers of the CNS 2017;14:1–8.

43. Liu P, Owashi K, Monnier H, Metanbou S, Capel C, Balédent O. Validating the accuracy of real-time phase-contrast MRI and quantifying the effects of free breathing on cerebrospinal fluid dynamics. Fluids and Barriers of the CNS 2024;21(1):25.

44. Gutiérrez-Montes C, Coenen W, Vidorreta M, Sincomb S, Martínez-Bazán C, Sánchez A, Haughton V. Effect of normal breathing on the movement of CSF in the spinal subarachnoid space. American Journal of Neuroradiology 2022;43(9):1369–1374.

45. Kollmeier JM, Gürbüz-Reiss L, Sahoo P, Badura S, Ellebracht B, Keck M, Gärtner J, Ludwig H-C, Frahm J, Dreha-Kulaczewski S. Deep breathing couples CSF and venous flow dynamics. Scientific reports 2022;12(1):2568.

46. Vikner T, Johnson KM, Cadman RV, Betthauser TJ, Wilson RE, Chin N, Eisenmenger LB, Johnson SC, Rivera-Rivera LA. CSF dynamics throughout the ventricular system using 4D flow MRI: associations to arterial pulsatility, ventricular volumes, and age. Fluids and Barriers of the CNS 2024;21(1):68.

47. Kaipainen AL, Martoma E, Puustinen T, Tervonen J, Jyrkkänen H-K, Paterno JJ, Kotkansalo A, Rantala S, Vanhanen U, Leinonen V. Cerebrospinal fluid dynamics in idiopathic intracranial hypertension: a literature review and validation of contemporary findings. Acta Neurochir (Wien) 2021;163(12):3353–3368.

48. Sagirov AF, Sergeev TV, Shabrov AV, Yurov AYe, Guseva NL, Agapova EA. Postural influence on intracranial fluid dynamics: An overview. J Physiol Anthropol 2023;42(1):5.

49. Heiss JD, Patronas N, DeVroom HL, Shawker T, Ennis R, Kammerer W, Eidsath A, Talbot T, Morris J, Eskioglu E. Elucidating the pathophysiology of syringomyelia. J Neurosurg 1999;91(4):553–562.

50. Gao JH, Holland SK, Gore JC. Nuclear magnetic resonance signal from flowing nuclei in rapid imaging using gradient echoes. Medical physics 1988;15(6):809–814.

51. Diorio TC, Nair VV, Patel NM, Hedges LE, Rayz VL, Tong Y. Real-time quantification of in vivo cerebrospinal fluid velocity using the functional magnetic resonance imaging inflow effect. NMR Biomed 2024:e5200.

52. Roefs EC, Eiling I, de Bresser J, van Osch MJ, Hirschler L. BOLD-CSF dynamics assessed using real-time phase contrast CSF flow interleaved with cortical BOLD MRI. Fluids and Barriers of the CNS 2024;21:107.

53. Ashenagar B, Gomez DE, Lewis LD. Modeling dynamic inflow effects in fMRI to quantify cerebrospinal fluid flow. Imaging Neuroscience 2025.

54. Beall EB, Lowe MJ. SimPACE: generating simulated motion corrupted BOLD data with synthetic-navigated acquisition for the development and evaluation of SLOMOCO: a new, highly effective slicewise motion correction. Neuroimage 2014;101:21–34.

55. Verma T, Cohen-Adad J. Effect of respiration on the B0 field in the human spinal cord at 3T. Magnetic resonance in medicine 2014;72(6):1629–1636.

56. Qvarlander S, Sundström N, Malm J, Eklund A. CSF formation rate—a potential glymphatic flow parameter in hydrocephalus? Fluids and Barriers of the CNS 2024;21(1):55.

57. Edsbagge M, Tisell M, Jacobsson L, Wikkelso C. Spinal CSF absorption in healthy individuals. American Journal of Physiology-Regulatory, Integrative and Comparative Physiology 2004;287(6):R1450–R1455.

